# Elevated CO_2_ induces age-dependent restoration of growth and metabolism in gibberellin-deficient plants

**DOI:** 10.1101/555292

**Authors:** Karla Gasparini, Lucas C. Costa, Fred A. L. Brito, Thaline M. Pimenta, Flávio Barcellos Cardoso, Wagner L. Araújo, Agustín Zsögön, Dimas M. Ribeiro

## Abstract

*Main conclusion* The effect of elevated [CO_2_] on the growth of tomato plants with reduced GA content is influenced by developmental stage.

The increase of carbon dioxide (CO_2_) in the atmosphere during the last decades has aroused interest in the function of this gas in the growth and development of plants. Despite the known association between elevated CO_2_ concentration ([CO_2_]) and plant growth, its effects in association with gibberellin (GA), plant hormone that regulates de major aspects of plant growth, are still poorly understood. Therefore, we evaluated the effect of elevated [CO_2_] on growth and primary metabolism in tomato plants with drastic reduction in GA content (*gib-1*) at two different growth stages (21 and 35 days after germination, dag). Disruption on growth, photosynthetic parameters and primary metabolism were restored when *gib-1* plants were transferred to elevated [CO_2_] at 21 dag. Elevated [CO_2_] also stimulated growth and photosynthetic parameters in Wild type (WT) plants at 21 dag, however, minor changes were observed in the level of primary metabolites. At 35 dag, elevated [CO_2_] did not stimulate growth in WT plants and *gib-1* mutants showed their characteristic stunted growth phenotype.

## Introduction

Plant growth and development involve many endogenous and environmental signals that interact with the plant’s genetic program to determine plant architecture (Wang and Li 2008; Achard et al. 2009). Fundamental to this process are chemical regulators known as plant hormones (Santner et al. 2009). Among them, gibberellins (GAs) regulate major aspects of plant growth and development, including seed germination, stem elongation, leaf expansion, trichome development and flowering induction (Hedden and Thomas 2016). Many of these aspects are controlled by the capacity of GAs to stimulate cell division and elongation (de Lucas et al. 2008; Achard et al. 2009), through degradation of transcriptional repressor DELLA proteins (Dill et al. 2001; Alvey and Harberd 2005).

GAs are synthesized via terpenoid pathway by the action of terpene synthase, cytochrome P450 oxygenase and 2-oxoglutarate-dependent dioxygenases (2-ODDs) in plastids, the endomembrane system and the cytosol, respectively. 2-ODDs are 2-oxoglutarate dependent, the key intermediate of one the most fundamental biochemical pathways in carbon metabolism -the tricarboxylic acid (TCA) cycle- and a point of connection between carbon and nitrogen metabolism (Araújo et al. 2014). In this way, the dependence on 2-oxoglutarate (2-OG) links GA biosynthesis and homeostasis directly with primary metabolism (Lancien et al. 2000), since most GA oxidases (GA_ox_) are 2-OG dehydrogenase (2-ODDs) (GA_3ox_, GA_20ox_, GA_2ox_).

Alterations in ambient [CO_2_] expected for the next few years (IPCC, 2014) may impact plant growth and development (Kimball 2016). Succinctly, [CO_2_] directly influences the rate of CO_2_ assimilation by Rubisco, and consequently gas exchange rates, which could influence plant growth and crop productivity (Campbell et al. 1988; Igamberdiev 2015; Galmés et al. 2017). Particularly in C3 plants, CO_2_ is a limiting substrate for photosynthesis, and elevated [CO_2_] usually leads to an increase in photosynthetic assimilation rates and a decrease in photorespiration, stimulating production of sugars and biomass accumulation (Ainsworth and Rogers 2007; Ainsworth 2008; Högy et al. 2010). Higher sugar availability is a trigger for plant growth at elevated [CO_2_] (Taylor et al. 1994; Masle 2000; Ferris et al. 2001). However, it is the sink capacity of the plant to use or store additional photoassimilate that determines photosynthesis stimulation and growth at elevated [CO_2_] (Arp 1991). In soybean (*Glycine max*), for instance, growth habit was a determinant factor of photosynthetic acclimation at elevated [CO_2_] (Ainsworth et al. 2004). In this case, the inability to form sufficient sinks in determinate-growth plants contributed to feedback photosynthesis acclimation, suggesting the participation of sugar sensing and signaling in the growth responses (Paparelli et al. 2013; Wang and Ruan 2013; Lastdrager et al. 2014).

Plant hormones act as chemical mediators to control growth and development in response to elevated [CO_2_]. Increase in GAs in response to elevated [CO_2_] have consistently been reported in several species such as *Ginkgo biloba* L.(Li et al. 2002), *Arabidopsis* (Teng et al. 2006) and *Populus* (Liu et al. 2014). Furthermore, growth reduction of *Arabidopsis* treated with the GA biosynthesis inhibitor paclobutrazol (PAC) was reverted by elevated [CO_2_] (Ribeiro et al. 2012). This suggests that plant growth at elevated [CO_2_] may be partially coupled with the effects of GA (Ribeiro et al. 2012), and that elevated [CO_2_] and GA act could in similar pathways related with plant growth. Despite this circumstantial evidence of association between elevated [CO_2_] and plant hormones, little is known about how elevated [CO_2_] coordinates plant growth together with GA.

Tomato (*Solanum lycopersicum* L.) is one of the most important horticultural crops in the world and has been widely used as a model organism in several fields of plant research (Kimura and Sinha 2008). The availability of monogenic mutant collections represents a powerful tool for the study of gene function and ecophysiological interactions (Carvalho et al. 2011). For example, three mutants in GA biosynthesis (*gibberellin deficient 1, 2 and 3, gib-1, gib-2* and *gib-3* respectively) were identified in tomato plants (Koornneef et al. 1990). The *gib-1* mutant shows reduction in ent-copalyl diphosphate synthase activity, the first enzyme involved in GA biosynthesis leading to a dwarf phenotype due to the drastic reduction in GA content (Bensen and Zeevaart 1990). Different to *gib-1*, both *gib-2* and *gib-3* show less conspicuous reductions in growth (Koornneef et al. 1990). Thus, the availability of these GA-related mutants makes tomato plants a useful model for the study of combinatorial effects of reduced GA content and elevated [CO_2_] on the plant growth.

Since elevated [CO_2_] influenced growth and metabolism of *Arabidopsis* treated with PAC (Ribeiro et al. 2012), here we investigated growth and metabolic responses in tomato plants with drastic reduction in GA content (*gib-1*) transferred to elevated [CO_2_] at two different growth stages (21 and 35 days after germination, dag). Mutant *gib-1* plants cultivated in ambient [CO_2_] showed stunted growth and reduced biomass accumulation, alterations in photosynthetic parameters and disruption in primary metabolism. Transfer to elevated [CO_2_] stimulated growth and most of primary metabolism of GA-deficient plants at 21 dag, but not 35 dag. We discuss the influence of elevated [CO_2_] on the growth of tomato plants with reduced GA content and how growth can be influenced by developmental stage in tomato plants submitted to elevated [CO_2_].

## Material and methods

### Growth conditions and experimental design

Seeds of tomato (*Solanum lycopersicum* L.) cv. Moneymaker and GA deficient *gib-1* mutants (kindly donated by M. Koornneef, Max Planck Institute for Plant breeding Research, Cologne, Germany) were germinated in Petri dish containing two layers of filter paper soaker in distilled water. After germination, the seedlings were transferred to pots (1,7L) containing commercial substrate (Tropstrato HT Hortaliças, Vida Verde), supplemented with NPK 20:5:20 fertilizer and cultivated as previously described in Vicente et al. (2015). Twenty-one and 35 days after germination, tomato plants were transferred to open top chamber under either ambient (400 μmol mol^-1^) and elevated (750 μmol mol^-1^) [CO_2_]. Plants were maintained inside the open top chambers for 21 days. The experiment was conducted in greenhouse localized at Universidade Federal de Viçosa (20° 45’S, 42° 15’W).

### Growth analysis

Stem length was measured every two days. At the end of the 21-day period at [CO_2_] treatment, plants were harvested and divided into leaves, stem and roots. Leaf area was measured using a planimeter (Li-Cor Model 3100 Area Meter, Lincoln, NE, USA). Shoot and root biomass were measured from the dry weight of leaves, stem and roots. Relative growth ratio (RGR) and specific leaf area (SLA) were determined as described by Hunt (1982).

### Leaf anatomy

Leaf discs were collected from the center of the third leaf and fixed in FAA50 (Formaldehyde, acetic acid and ethanol 50%) for 48 h, and then stored in ethanol 70% according to Johansen (1940). Then the plant material was dehydrated in ethanolic series and included in methacrylate (Historesin-Leica), according to the manufacturer’s recommendations. For light microscope observation (AX-70 TRF, Olympus Optical, Tokyo, Japan), cross sections 5 µm thick were obtained with an automatic advanced rotary microtome (model RM2155, Leica microsystems Inc., Deerfield, USA), were stained with toluidine blue then photographed using a digital camera (Zeiss AxioCam HRc, Göttinger, Germany). Anatomical features, as leaf thickness and thickness cell layers, were evaluated using Image J (NIH, Bethesda, Ma).

### Gas exchanges and fluorescence measurements

The net rate of carbon assimilation (*A*), stomatal conductance (g_s_), internal CO_2_ concentration (C_i_) and fluorescence parameters were measured in third or fourth fully-expanded leaf from the botton, using infrared gas analyzer (Li 6400XT, Li-Cor, Lincoln, USA) equipped with integrated fluorescence chamber head (Li-6400-40; Li-Cor Inc.). The measurements were conducted with [CO_2_] supply of 400 and 750 μmol CO_2_ mol^-1^ air with artificial photosynthetically active radiation level of 500 μmol of photons m^-2^ s^-1^, matching the greenhouse irradiance value. The rate of mitochondrial respiration in darkness (*R*_D_) was measured on dark-adapted leaves for at least 2 hours after the end of the light period. Using the values of these parameter, mitochondrial respiration in light (*R*_L_) was estimated according to Lloyd et al. (1995) as *R*_L_ =[0.5 – 0.05ln(PPFD)]×*R*_D_. The photorespiratory rate of Rubisco (*R*_P_) was estimated according Valentini et al. (1995), as *R*_P_=1/12[ETR – 4(*A*+*R*_L_)]. *A*/C_i_ curves were measured at saturating irradiance (1000 μmol photons m^-2^ s^-1^) as described Barbosa et al. (2018). From these curves, maximum rate of carboxylation (*V*_cmax_) and the maximum rate of electron transport (*J*_max_) were calculated as proposed by Sharkey et al. (2007).

### Determination of metabolic levels

Samples of third leaf, midpoint region of stem and root were collected, immediately frozen in liquid nitrogen, and stored at −80°C until further analysis. Glucose, fructose, sucrose and starch levels were determined as described by Fernie et al. (2001). Amino acids levels were determined as described by Gibon et al. (2004). Protein levels were measured as described by Bradford (1976), with modifications (Gibon et al. 2004). The levels of nitrate were measured as described by Fritz et al. (2006). Chlorophylls were extracted using acetone 80% and the content of them was determined as described by Lichtenthaler (1987). The levels of all others metabolites were quantified by gas chromatography mass spectrometry (GC-MS) following the protocol described by Lisec et al. (2006).

### Statistical analysis

The experiments were designed in a completely randomized distribution. Differences described are systematically statistically grounded based on ANOVA, where P<0.05 was considered significant. If ANOVA showed significant effects, Student’s t test (P<0.05) was used to determine differences between each treatment and control. All statistical analyses were made using Statistical Package for the Social Sciences for Windows statistical software (SPSS).

## Results

### Elevated [CO_2_] stimulates impaired growth of the gib-1 mutant

The *gib-1* tomato mutant shows impaired growth due to reduction in the activity of ent-copalyl diphosphate synthase, a key enzyme in the GA biosynthesis pathway (Bensen and Zeevaart 1990). Under ambient [CO_2_], *gib-1* plants showed severe reductions in shoot (−82%) and root (−29%) biomass, as well as leaf area (−80%) and stem length (−77%), compared with WT plants (Fig. 1). In addition, SLA and RGR were lower in *gib-1* mutants, compared with WT plants at ambient [CO_2_] (Supplementary Figure 1). Elevated [CO_2_] did not affect shoot and root biomass of WT plants but led to increased leaf area (22%) and stem length (41%), compared with WT plants at ambient [CO_2_] (Fig. 1). In the *gib-1* mutant, elevated [CO_2_] restored shoot and root biomass, as well as leaf area, to similar values as WT plants at ambient [CO_2_] (Fig. 1). Elevated [CO_2_] doubled the stem length in the *gib-1* mutant, compared to *gib-1* at ambient [CO_2_], but still fell short of the stem length of WT plants at ambient [CO_2_] (Fig. 1 d). *gib-1* plants in elevated CO_2_ also showed SLA and RGR values similar to those of ambient [CO_2_] WT plants (Supplementary Figure 1).

**Fig. 1.**
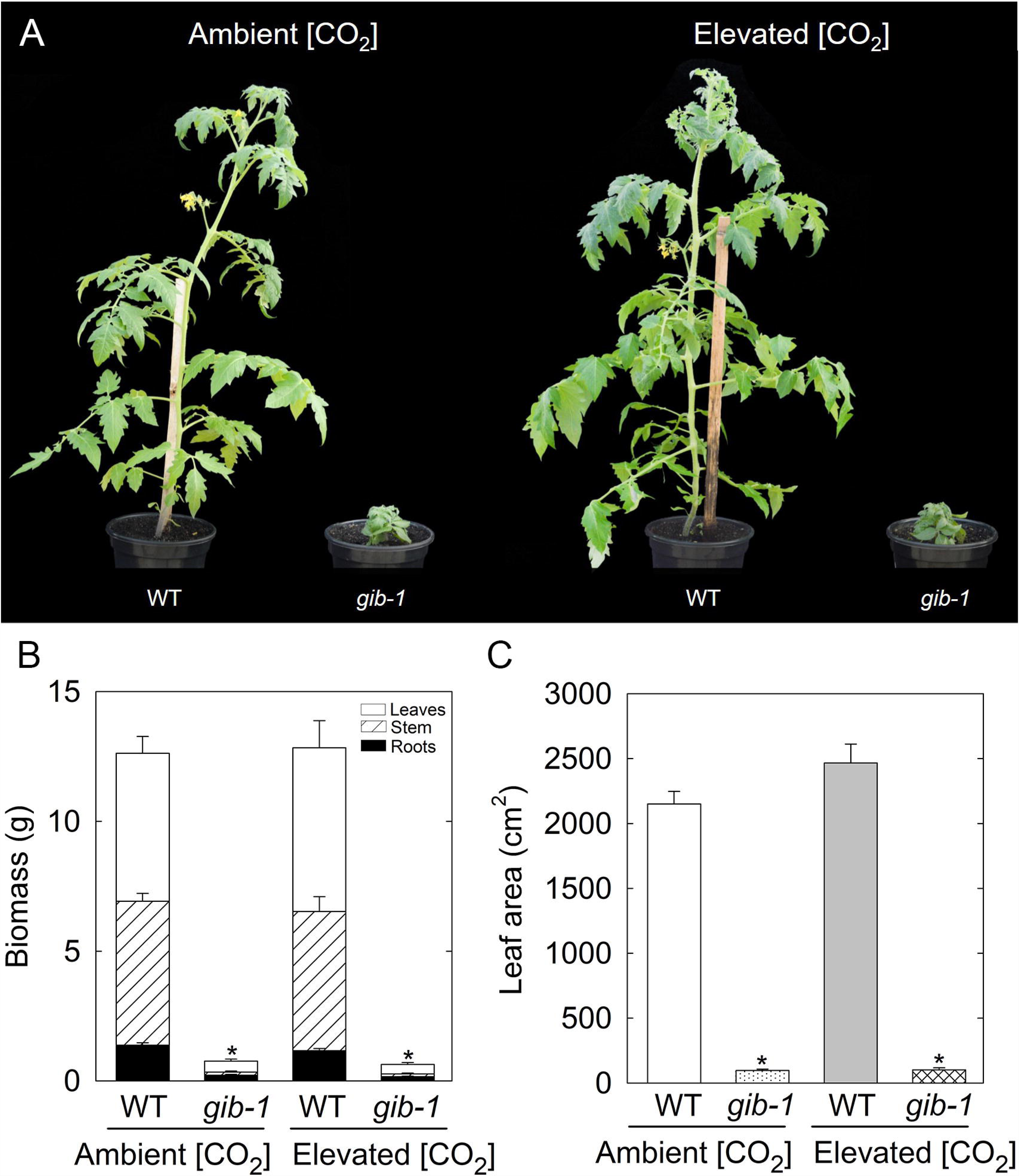
Effects of elevated [CO_2_] on growth of wild type (WT) and *gib-1* mutant 21 dag. a, phenotypes of WT and *gib-1* mutant grown at ambient and elevated [CO_2_]. b, shoot and root biomass. c, leaf area. d, stem length. Measurements were done in tomato plants after 21 days of growing at 400 or 750 μmol CO_2_ mol^-1^. Asterisks indicate values determined by Student’s t test to be significantly different from WT plants in ambient [CO_2_] (P<0.05). Values are means ± standard error of 10 replicates.

### Elevated [CO_2_] restores leaf anatomy in gib-1 plants

Given the stimulatory effects of GAs and elevated [CO_2_] on cell expansion and division (Taylor et al. 1994, 2005; Achard et al. 2009) we decided to analyze how elevated [CO_2_] influences leaf anatomy of *gib-1* mutant and WT plants. Under ambient [CO_2_], leaf cross-sections revealed considerable visual differences between *gib-1* mutants and WT plants (Fig. 2 a and b). Leaf thickness was increased (35%) in *gib-1* compared to WT due to a 18% and 62% increase in the thickness of palisade and spongy parenchyma, respectively (Fig. 2 e, f and g). The greater thickness of spongy parenchyma resulted in the reduction of the palisade-to-spongy parenchyma ratio in *gib-1* (Fig. 2 h)

**Fig. 2.**
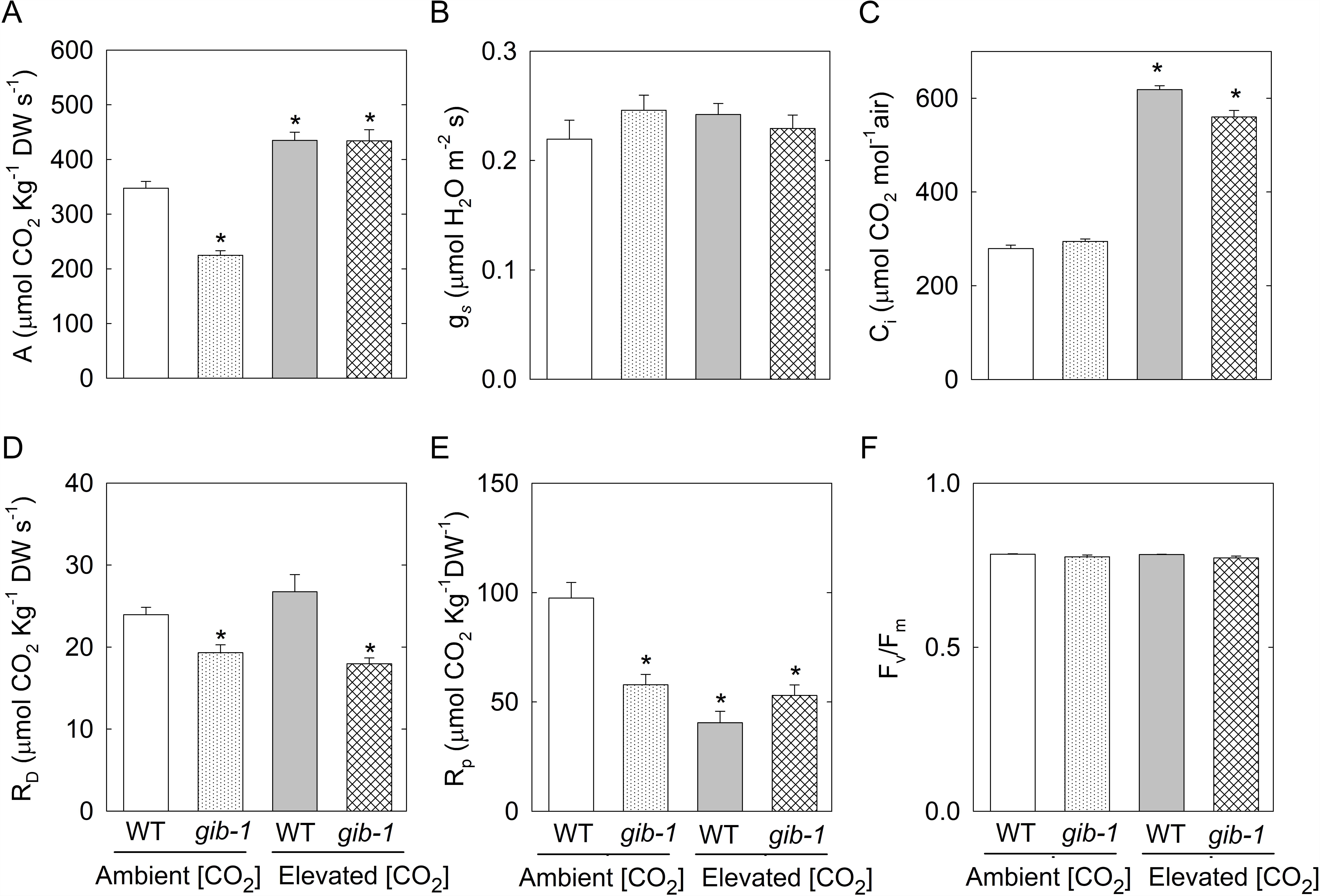
Effects of GA and elevated [CO_2_] at leaf anatomy of wild type (WT) and *gib-1* mutant at 400 or 750 μmol CO_2_ mol^-1^. a, b, c and d cross section of the third leaf of the wild type and *gib-1* plants at 400 or 750 μmol CO_2_ mol^-1^ (Scale bar: 100μm). e, total leaf thickness. f, palisade parenchyma thickness. g, spongy parenchyma thickness. h,p alisade: spongy parenchyma ratio. i, upper epidermis thickness. j, lower epidermis thickness. Asterisks indicate values determined by Student’s t test to be significantly different from WT plants at ambient [CO_2_] (P<0.05). Values are means ± standard error of 6 replicates. UE, Upper epidermis; LE, lower epidermis; PP, palisade parenchyma; SP, spongy parenchyma.

Under elevated [CO_2_], *gib-1* leaf cross-sectional appearance was very similar to that of WT plants at ambient [CO_2_] (Fig. 2 a and d). At elevated [CO_2_], total leaf thickness, palisade and spongy parenchyma thickness, and the palisade-to-spongy parenchyma ratio of *gib-1* plants were similar to those of WT plants at ambient [CO_2_] (Fig. 2 e, f, g and h). Thickness of either upper or lower epidermis was not altered in *gib-1* between [CO_2_] levels (Fig. 2 i and j). As for WT plants, most leaf anatomical parameters were similar between treatments, except for upper epidermis thickness, which was reduced in elevated [CO_2_] plants compared to ambient [CO_2_] (Fig. 2).

### Elevated [CO_2_] restore photosynthetic function in the gib-1 mutant

We next investigated the combined effects of reduction in endogenous GA content and elevated [CO_2_] on gas exchange in *gib-1* mutant tomato plants. Under ambient [CO_2_], the *gib-1* mutant showed a marked reduction (∼45%) in *A*, compared with WT plants (Fig. 3 a). In addition, the *gib-1* mutation led to reductions in *R*_D_ (−52%), *V*_cmax_ (−45%), *J*_max_ (−37%), *R*_P_ (−41%) and ETR (−44%) at ambient [CO_2_]. Under ambient [CO_2_], C_i_ was unaffected in the *gib-1* mutant, compared with WT plants (Fig. 3)

**Fig. 3.**
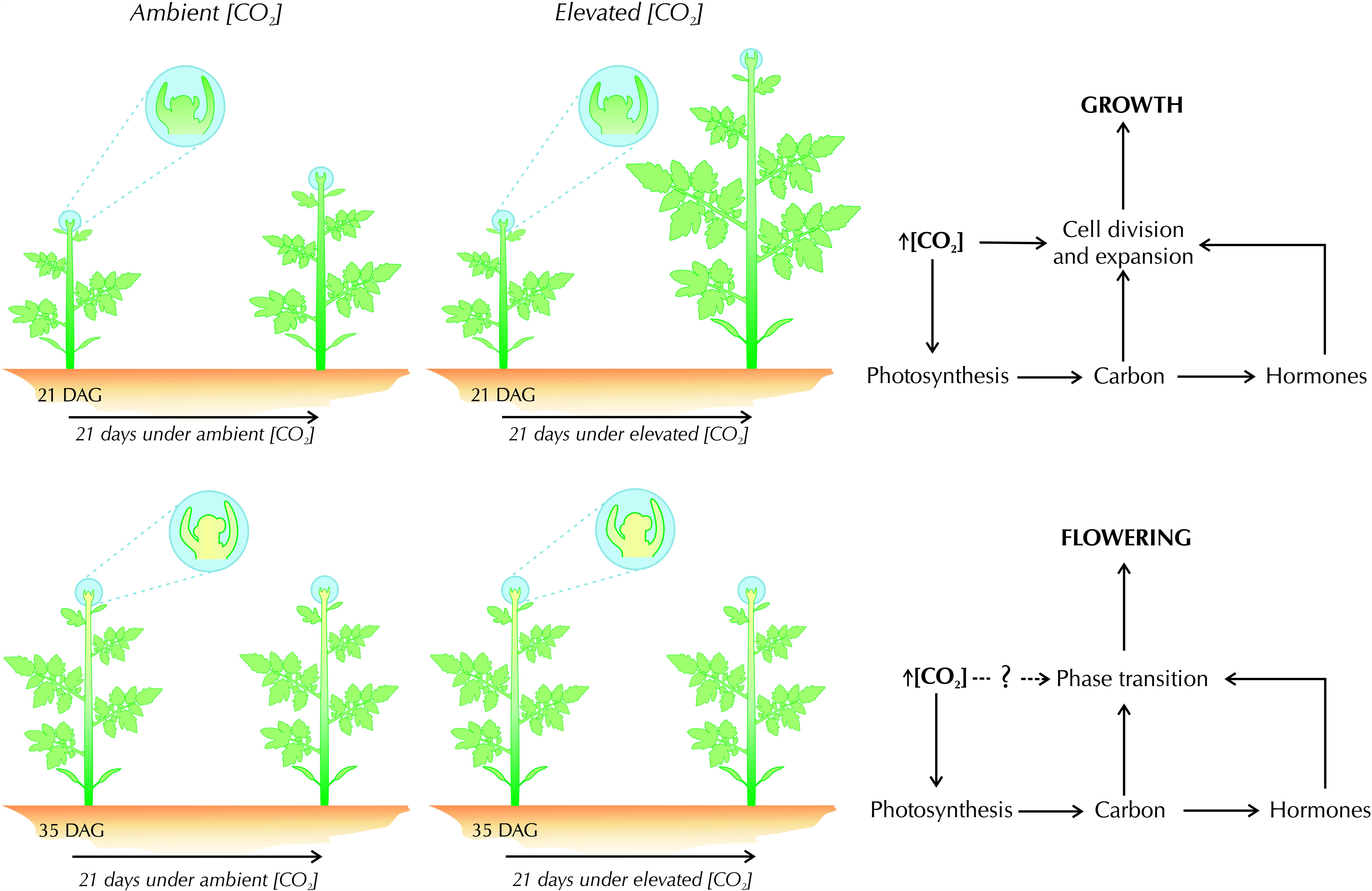
Changes in gas exchange and fluorescence parameters in wild type (WT) and *gib-1* mutant at 400 or 750 μmol CO_2_ mol^-1^. a, net rate of carbon assimilation (*A*). b, rate of mitochondrial respiration in darkness (*R*_D_). c, maximum rate of carboxylation (*V*_cmax_). d, maximum rate of electron transport (*J*_max_). e, photorespiratory rate of Rubisco (*R*_P_). f, electron transport rate (ETR). g, internal CO_2_ concentration (C_i_). h, stomatal conductance (g_s_). i, variable to maximum fluorescence ratio (*F*_v_/*F*_m_). Asterisks indicate values determined by Student’s t test to be significantly different from WT plants in ambient [CO_2_] (P<0.05). Values are means ± standard error of 10 replicates.

Under elevated [CO_2_], *A, R*_D_ and C_i_ values were higher in both WT and *gib-1* compared to ambient [CO_2_] WT plants (Fig. 3 a, b and g). Elevated [CO_2_] increased *J*_max_ in *gib-1* but not in WT plants, compared to ambient [CO_2_] WT plants (Fig. 3 d). *R*p was considerably reduced in both genotypes at elevated [CO_2_] (Fig. 3 e). Interestingly, *g*_s_ remained stable in WT and *gib-1* under both ambient and elevated [CO_2_] (Fig. 3 h). The *F*v/Fm ratio was close to ideal values for non-stressed leaves (∼0.83) in WT plants and *gib-1* mutant in both [CO_2_] regimes (Fig. 3 i)

### Nitrogen metabolism is altered by elevated [CO_2_] in gib-1 mutant plants

Elevated [CO_2_] typically reduces nitrogen content in the tissues of some plant species (Bloom et al. 2002, 2010, 2014; Taub and Wang 2008). We thus determined the content of some of the main nitrogenous compounds in leaves, stem and roots of WT and *gib-1* plants under ambient and elevated [CO_2_]. Nitrate levels remained generally unchanged across treatments, except for an increase in *gib-1* at ambient [CO_2_] (Supplementary Fig. 2). Alterations in amino acids levels were found for the *gib-1* mutant at ambient [CO_2_]. Whereas amino acids levels were reduced in leaf (35%), an increase was observed in the stem (76%) and roots (41%) (Supplementary Fig. 2). At elevated [CO_2_], amino acids levels remained unaltered in WT plants and *gib-1* mutants, compared with WT plants at ambient [CO_2_]. At ambient [CO_2_], leaf protein content was reduced by 24%, while it increased approximately four-fold in stems of *gib-1* mutants (Supplementary Fig. 2). Leaf protein was lower in *gib-1* mutant and WT plants under elevated [CO_2_] (Supplementary Fig. 2). In the roots, protein levels were increased in both *gib-1* and WT plants at elevated [CO_2_] (Supplementary Fig. 2). Under ambient [CO_2_], chlorophyll analysis of *gib-1* mutants showed increases of 18% and 65% in the leaf and stem, respectively. Elevated [CO_2_] did not affect leaf chlorophyll content in WT plants.

### Changes in carbohydrate content and partitioning associated with GA content and elevated [CO_2_]

Photosynthetic assimilation of CO_2_ and GA are coupled to carbon metabolism and consequently to carbohydrates content (Ainsworth et al. 2002; Ainsworth and Long 2005; Leakey et al. 2009; Paparelli et al. 2013). Since carbohydrates are essential to the fundamental process required to plant growth (Eveland and Jackson 2012), we investigated the levels of soluble sugars and starch in WT and *gib-1* mutant grown both at ambient and elevated [CO_2_]. Under ambient [CO_2_], *gib-1* mutants showed reduced levels of leaf and stem glucose (Supplementary Fig. 3) compared with WT plants at ambient [CO_2_]. By contrast, glucose level in roots increased in *gib-1* mutants at ambient [CO_2_] (Supplementary Fig. 3). Under ambient [CO_2_], the reduction of leaf fructose in *gib-1* mutants was accompanied by an increase in stem and roots in these plants, compared with WT plants at ambient [CO_2_]. The levels of sucrose showed a similar pattern as those of fructose in *gib-1* plants grown at ambient [CO_2_]. Growth of *gib-1* mutant at ambient [CO_2_] resulted in reduced leaf starch.

Elevated [CO_2_] did not affect glucose levels in WT plants but restored them in leaves of *gib-1* mutants. Although higher than *gib-1* at ambient [CO_2_], the level of glucose in stem was reduced in *gib-1* mutants grown at elevated [CO_2_]. Moreover, the level of glucose in the root increased in *gib-1* mutant at elevated [CO_2_]. Although elevated [CO_2_] did not affect fructose content in leaf, it increased it in stem and roots of WT plants, compared with WT plants at ambient [CO_2_]. At elevated [CO_2_], *gib-1* mutants showed increased fructose content in all plant organs analyzed, compared with WT plants at ambient [CO_2_]. Elevated [CO_2_] did not affect sucrose content in WT plants but restored it in the *gib-1* mutant. The level of starch increased in WT plants and *gib-1* mutants under elevated [CO_2_], compared with WT plants at ambient [CO_2_] (Supplementary Fig. 3).

### Elevated [CO_2_] modifies part of the metabolic profile in gib-1 mutant

To investigate how the metabolism of each organ was modified by GA and elevated [CO_2_], we built a metabolic profile in WT and *gib-1* plants under ambient and elevated [CO_2_] (Fig. 4).

**Fig. 4.**
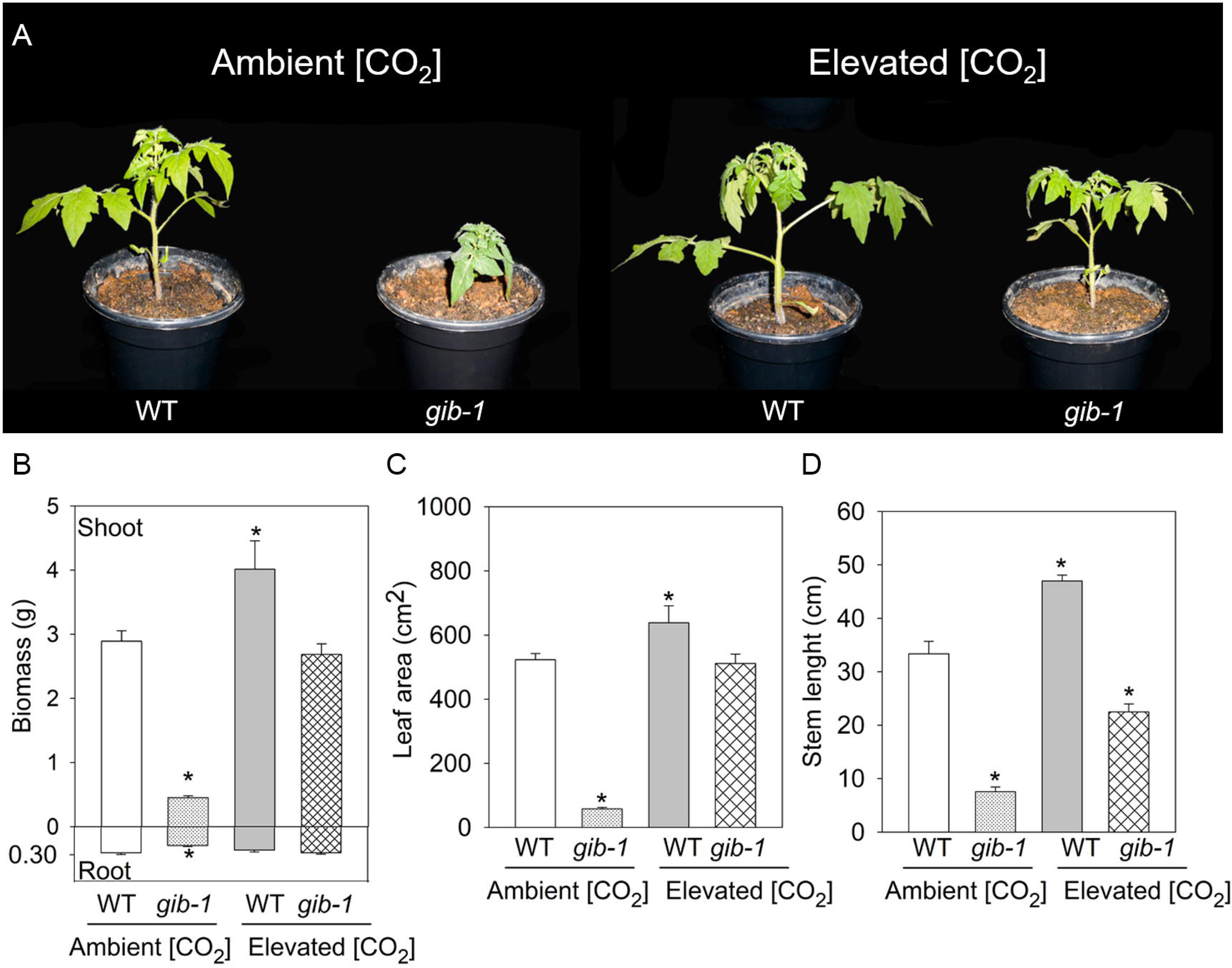
Relative metabolite level of leaf, stem and root from wild type (WT) and *gib-1* mutant at 400 or 750 μmol CO_2_ mol^-1^. Samples of leaf, stem and root were collected at the end of the light period from plants growing for 21 days at ambient or elevated [CO_2_]. Asterisks indicate values determined by Student’s t test to be significantly different from wild type (WT) plants in ambient [CO_2_] (P<0.05). Values are means ± standard error of 6 replicates. Data are normalized with respect to mean response calculated for the wild type (WT) plants in ambient [CO_2_]. nd, not detected. The full dataset from the metabolite profiling study is available as Supplementary Table S1.

Under ambient [CO_2_], the levels of 34 (out of 40) leaf metabolites were affected in the *gib-1* mutant, of which 30 were reduced and 4 increased compared with WT plants at ambient [CO_2_] (Fig. 4). Among the metabolites whose level increased are 3-PGA, trehalose, oxalic acid and leucine. Elevated [CO_2_] affected few leaf metabolites in WT plants and restored the levels of 26 of the 34 metabolites affected in the *gib-1* mutant grown at ambient [CO_2_]. The levels of galactinol, glyceric acid, glycerol, mannose, 2-OG and proline were reduced in leaf both WT plants and *gib-1* mutants at elevated [CO_2_], while citric acid was increased. Elevated [CO_2_] increased the level of leaf GABA only in the *gib-1* mutant. Myo-inositol, lactate, glycine, ornithine, serine and valine remained unchanged in WT plants and *gib-1* mutant under both ambient and elevated [CO_2_].

Thirty-three metabolites were detected in the stem by GC-MS. In *gib-1* mutant grown at ambient [CO_2_], 13 metabolites were reduced, while 11 were increased, compared with WT plants at ambient [CO_2_]. Most metabolites were not affected by elevated [CO_2_] in the stem of WT plants, compared with WT plants at ambient [CO_2_]. However, the level of galactinol, mannose, malate and serine showed reduced and fructose increased levels in WT plants grown at elevated [CO_2_] in relation to WT plants at elevated [CO_2_]. Elevated [CO_2_] restored 19 of 24 metabolites affected in stems of *gib-1* mutants at ambient [CO_2_]. The level of fructose, asparagine, glutamate, glutamine remained increased, while malate and serine were reduced in *gib-1* mutants at elevated [CO_2_], compared with WT at ambient [CO_2_]. No change was observed in myo-inositol, lactate, pyruvate, GABA, glycine, leucine, valine and β-alanine (Fig. 4).

In roots, most amino acids were increased in *gib-1* mutants at ambient [CO_2_]. In addition, the level of fructose, glucose, 3-PGA, sucrose and 2-OG also increased in the root of *gib-1* mutants, compared with WT at ambient [CO_2_]. The level of fumarate, malate and oxalacetate decreased in *gib-1* mutants grown at ambient [CO_2_]. Elevated [CO_2_] led to increase in the level of fructose, galactinol, 3-PGA, succinate, glycine, phenylalanine, tyrosine and β-alanine in WT plants and *gib-1* mutant, while reducing the level of malate in these plants compared with WT plants at ambient [CO_2_]. At elevated [CO_2_], an increase in glucose, citrate, isocitrate, glutamate and serine was observed only in the roots of *gib-1* mutant, compared with WT plants at ambient [CO_2_]. Lastly, no changes were observed in myo-inositol, aconitate, lactate, pyruvate, GABA, ornithine and proline in the roots (Fig. 4).

### Elevated [CO_2_] stimulates growth in tomato plants 21 dag, but not 35 dag

The effect of elevated [CO_2_] on plant growth and development was described as age-dependent (Ainsworth 2008; Franks 2013). Thus, the influence of [CO_2_] on growth, gas exchange and primary metabolism was evaluated in WT and *gib-1* mutant submitted at 35 dag at ambient and elevated [CO_2_]. Under ambient [CO_2_], *gib-1* showed a drastic reduction in growth, biomass (∼93%) and leaf area (∼95%), compared to WT plants (Fig. 5). Growth of WT and *gib-1* at elevated [CO_2_] did not affect the growth parameters evaluated at 35 dag (Fig. 5 b and c).

**Fig. 5.**
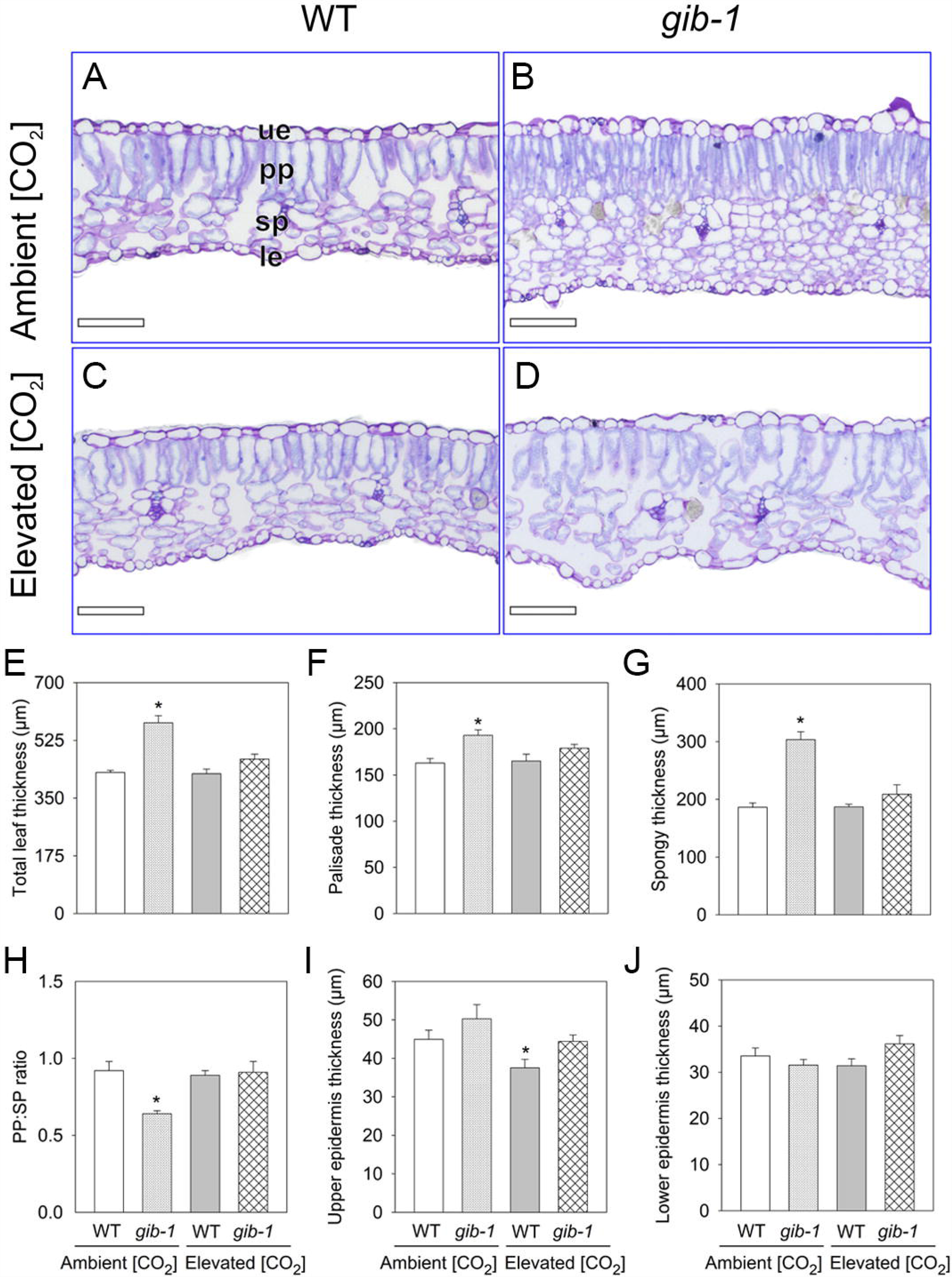
Effects of elevated [CO_2_] on growth of wild type (WT) and *gib-1* mutant 35 dag. a, phenotypes of WT and *gib-1* mutant grown at ambient and elevated [CO_2_]. b, total biomass. c, leaf area. Measurements were done in tomato plants after 21 days of growing at 400 or 750 μmol CO_2_ mol^-1^. Asterisks indicate values determined by Student’s t test to be significantly different from WT plants in ambient [CO_2_] (P<0.05). Values are means ± standard error of 10 replicates.

Under ambient [CO_2_], reduction GA content in *gib-1* mutant decreases *A* (∼35%), *R*_D_ (∼19%) and *R*_P_ (∼40%) at 35 dag, compared with WT (Fig 6 a, d and e). Growth in elevated [CO_2_] increase *A* and Ci, while reduced *R*_p_ in both, WT and *gib-1* (Fig. 6 a, c and e). No alterations in WT plants was observed in *R*_D_, however, these parameters were decreased in *gib-1* mutant at elevated [CO_2_], compared to WT plants under ambient [CO_2_]. *g*_s_ remained stable in WT and *gib-1* under both ambient and elevated [CO_2_] (Fig. 6 b). The *F*v/Fm ratio was close to ideal values for non-stressed leaves (∼0.83) (Fig. 6 f).

**Fig. 6.**
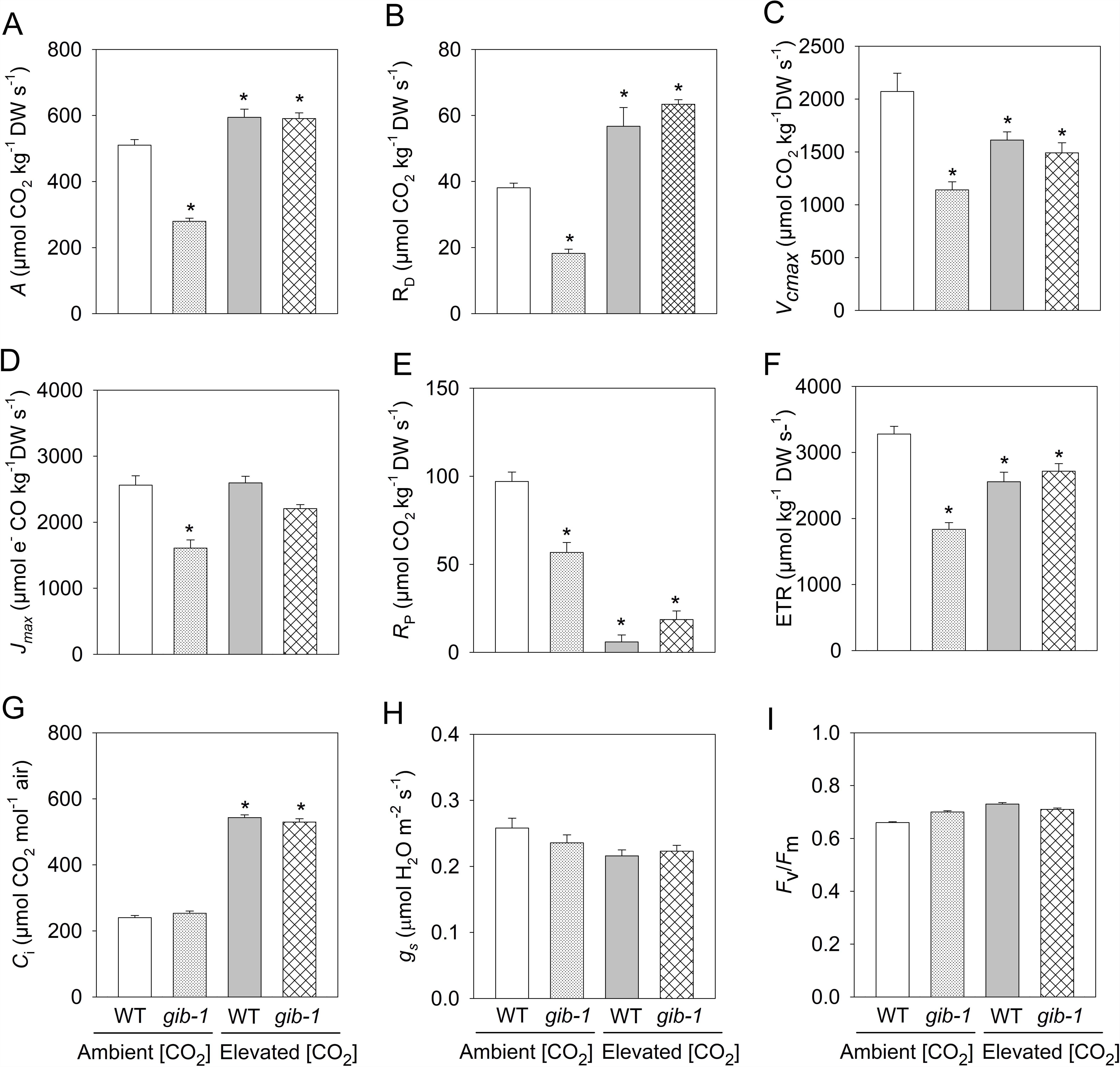
Changes in gas exchange in wild type (WT) and *gib-1* mutant at 400 or 750 μmol CO_2_ mol^-1^, 35 dag. a, net rate of carbon assimilation (*A*). b, stomatal conductance (g_s_). c, internal CO_2_ concentration (C_i_). d, rate of mitochondrial respiration in darkness (*R*_D_). e,p hotorespiratory rate of Rubisco (*R*_P_). f, variable to maximum fluorescence ratio (*F*_v_/*F*_m_). Asterisks indicate values determined by Student’s t test to be significantly different from WT plants in ambient [CO_2_] (P<0.05). Values are means ± standard error of 10 replicates.

Sugar evaluation of *gib-1* mutant submitted to ambient [CO_2_] at 35 dag shows reduction in leaf glucose (∼50%) and fructose (∼87%) and increased in sucrose compared to WT plants at ambient [CO_2_]. Under elevated [CO_2_] WT plants did not differ in leaf sugar content when compared to WT plants at ambient [CO_2_]. *gib-1* mutants kept reduction on glucose (∼51%), fructose (∼88%), and increase in sucrose (∼27%) at elevated [CO_2_] (Supplementary Fig. 4)

## Discussion

Elevated [CO_2_] stimulates growth at least in part in a GA-independent manner in *Arabidopsis* treated with GA synthesis inhibitor (PAC) (Ribeiro et al. 2012). Here, we evaluated the effect of [CO_2_] in tomato mutants with drastic reduction in GA content. GA regulates major aspects of growth and here we show that elevated [CO_2_] can directly stimulate mechanisms that compensate GA deficiency in *gib-1* mutants. However, this effect is strongly dependent on plant age.

### Source-sink relationships determines carbohydrate allocation

Alterations in endogenous level of GAs change the pattern of growth and biomass allocation (Nagel et al. 2001). The concentration of GAs in the *gib-1* mutant is insufficient to maintain normal leaf and stem growth, however, a less drastic effect is observed on root growth (Fig. 1). The marked reduction in growth and shoot biomass allocation impaired RGR in *gib-1* mutants (Supplementary Fig. 1). RGR has a positive relation with leaf mass ratio and SLA (Gleeson and Tilman 1992; Poorter and van der Werf 1998), parameters that are strongly affected by GA. Elevated [CO_2_] reestablished biomass allocation and SLA in *gib-1* mutants, which in turn influenced positively RGR.

Leaf anatomy influences photosynthetic capacity, determining the diffusion of CO_2_ through the mesophyll (Terashima et al. 2011; Tomás et al. 2013). Reduction in GA content acts as a factor disrupting growth and leaf development, since GA acts in the cell division, cell expansion and mesophyll organization (Jiang et al. 2012). Impaired leaf expansion in GA-deficient plants leads to increased number of cells per unit leaf area (Jiang et al. 2012), giving *gib-1* the appearance of a highly packed mesophyll (Fig. 2). In addition, increased mesophyll thickness influenced the reduction of SLA in *gib-1* mutant under ambient [CO_2_] (Supplementary Figure 1).

The increase in leaf thickness accompanied by the reduction in intercellular spaces in the *gib-1* mutant may have contributed to the reduction in *A*, since they restrict the diffusion of CO_2_ through the mesophyll and to the carboxylation site of Rubisco (Evans and Caemmerer 1996; Terashima et al. 2001). Furthermore, reduced lamina size and overlapping leaves may cause self-shading in *gib-1*, which also impairs photosynthetic capacity.

Although *gib-1* presented reduced *A* (Fig. 3), CO_2_ fixation probably exceeded the demand for growth, leading to the accumulation of carbohydrates in the stem and root (Supplementary Fig. 3). Stem storage of excess photoassimilate during periods of low sink strength buffers against source-sink changes during the different stages of growth (Slafer 2003). Imbalance between source and sink can lead to downregulation of photosynthesis due to accumulation of non-structural carbohydrates (such as soluble sugars and starch) in leaves (Stitt and Krapp 1999; Ainsworth and Bush 2011; Sugiura et al. 2017). Thus, the allocation of carbohydrates to the stem could be a way of delaying photosynthetic inhibition in *gib-1* mutants. In addition, the larger fraction of carbohydrate allocated to the root supports root respiration, which is less affected by GA deficiency than shoot respiration (Nagel and Lambers 2002).

In general, elevated [CO_2_] increases carbon assimilation and the availability of carbohydrates, which contribute to increased plant growth (Ainsworth and Long 2005; Teng et al. 2006; Li et al. 2013). However, it is the ability to grow and produce new sinks that determines photoassimilates consumption, whose accumulation could result in the inhibition of photosynthesis. Dark respiration (*R*_D_) is closely related to carbon balance and therefore the availability of carbohydrates can interfere in *R*_D_. Elevated [CO_2_] accelerate the accumulation of carbohydrates, which leads to transcriptional up-regulation of genes associated with respiration pathways and *R*_D_ stimulus (Li et al. 2008; Leakey et al. 2009; Markelz et al. 2014; Watanabe et al. 2014). Thus, the increase in *R*_D_ observed in WT and *gib-1* mutants can be attributed to the increase in *A* and consequently to the increase of carbohydrates under elevated [CO_2_] (Fig. 3 and Supplementary Fig. 3).

### Elevated [CO_2_] induces metabolic homeostasis in GA deficient plants

Plant growth is dependent on the interaction between carbon and nitrogen metabolism, which are linked by the tricarboxylic acid (TCA) cycle (Nunes-Nesi et al. 2010). Although the physiological function of genes regulated by GAs has been addressed (Yamaguchi 2008), studies on the effect of GAs on energy metabolism are scarce. We did not observe any drastic differences in metabolites levels between ambient and elevated CO_2_ treatments for WT plants (Fig. 4). Enhanced plant growth under elevated [CO_2_] does not induce a massive remodeling of metabolism. Optimization of carbon and nitrogen acquisition under such conditions appears to be dependent mostly on fine-tuning of specific points of the metabolic network. GA-deficient plants, on the other hand, show a general reduction in levels of sugars and amino acids in the leaf and an increase in roots under ambient [CO_2_] (Fig 4 and Supplementary Fig. 2, 3). This is probably a consequence of the alteration in carbon allocation in *gib-1*, whereby the mutation leads to increased root-to-shoot biomass ratio.

Elevated [CO_2_] restored the levels of most metabolites in *gib-1* to the level of WT (Fig. 4). Alterations in the TCA cycle have been shown to influence GA levels (Margaretha et al. 2009; Araújo et al. 2012), as multiple enzymes in the GA biosynthetic pathway are dependent on a TCA cycle intermediate, 2-oxoglutarate. On the other hand, it is not clear if the reverse is true, *i.e.* how does altered GA impact primary metabolism?

GABA levels increased only in the leaves of *gib-1* under elevated [CO_2_] (Fig. 4). Increases in GABA concentration occur in response to extreme conditions, like temperature, dehydration, salinity, oxygen stress (Kinnersley and Turano 2000; Bouché and Fromm 2004). GABA provides an alternative pathway for the conversion of alpha ketoglutarate to succinate in the TCA cycle, and compromising the enzymes of the TCA cycle involved in the steps up to the production of succinate alters GABA shunt activity (Lemaitre et al. 2007; Fait et al. 2008). Exogenous GABA application improved growth of *Zea mays, Stellaria longipes* and *Lemna*, possibly by inducing cell elongation and division or/and by maintaining metabolic balance within plant tissues (Kathiresan et al. 1998; Kinnersley and Lin 2000; Li et al. 2016). Thus, elevated [CO_2_] in plants with reduction in GA can influence GABA content and consequently carbon flux, ameliorating *gib-1* growth at elevated [CO_2_].

### The effect of elevated [CO_2_] on gib-1 is age-dependent

Both elevated [CO_2_] and GA influence the expression of genes related to loosening and rearrangement of the cell wall, besides controlling the rate of cellular proliferation (Vogler et al. 2003; Yang et al. 2004; Achard et al. 2009; Ribeiro et al. 2012). We showed that GA, an essential hormone for the normal growth and development of plants, is dispensable for growth in tomato plants under elevated [CO_2_]. Interestingly, we observed that the effects of elevated [CO_2_] on tomato growth is age-dependent, regardless of GA content (Fig. 5). These results indicate the existence of a “sensitive phase” in which elevated [CO_2_] is able to influence growth in plants. In addition to CO_2_, the action of other environmental factors is also restricted to the “sensitive phase” in tomato plants. Calvert (1957) showed that the influence of temperature on tomato flowering occurs in the first two or three weeks from the seedling emergence. This stage corresponds to our treatment of plants submitted to elevated [CO_2_] with 21dag.

In Arabidopsis the most pronounced growth under elevated [CO_2_] was observed during the vegetative stage (Watanabe et al. 2014). As observed in Arabidopsis, the sensitivity of tomato plants to elevated [CO_2_] may be linked to the juvenile phase. The efficiency of enzymes that promote cell expansion and, consequently, plant growth depends on factors controlled by the stage of development, which limits growth to specific periods (Sloan et al. 2009). Addition of new extracellular polymers and remodeling of existing components in the primary cell walls marks the exponential phase of cell expansion. This is followed by cell wall thickening and rigidification to create secondary cell walls that enhance structural integrity, but reduce cell wall extension (Hall and Ellis 2013). Furthermore, the transition from the juvenile to the adult phase can determine plant architecture and growth pattern (Huijser and Schmid 2011; Poethig 2013). Phase transition is preceded by a change in the competence of the shoot to respond to stimuli that induce reproductive development (Poethig 2013). The changes in meristem identity during phase transition are accompanied by genetic reprogramming that may trigger changes in leaf and stem morphology, as well as alteration in growth rate (Poethig 2010, 2013). With the development of floral organs most of the photoassimilates are destined for the production of flowers and fruits and for the maintenance of respiration of reproductive structures (Obeso 2002).

In conclusion, elevated [CO_2_] favors photosynthesis and carbohydrate production, regardless of plant age. However, we showed here that plant age can indirectly influence carbon partitioning via changes in source-sink relationships. These changes are mostly driven by the growth phase, either juvenile or adult, whereby vegetative or reproductive structures will be favored as strong sinks. Plant hormones can act as integrators of growth and development, and GAs, in particular, control cell division and expansion. We have shown here that increased growth under elevated [CO_2_] in tomato does not require a functional gibberellin biosynthetic pathway. In the juvenile phase, gibberellin-deficient mutants can grow to the same extent as wild-type plants. In the adult phase, however, elevated [CO_2_] does not stimulate growth and gibberellin mutants show their characteristic stunted growth phenotype. This suggests that growth stimulation by [CO_2_] is highly dependent on plant developmental stage, possibly linked to the juvenile-to-adult phase transition (Fig. 7). Further work should explore the potential role of other hormones mediating growth stimulation by elevated [CO_2_].

**Fig. 7.**
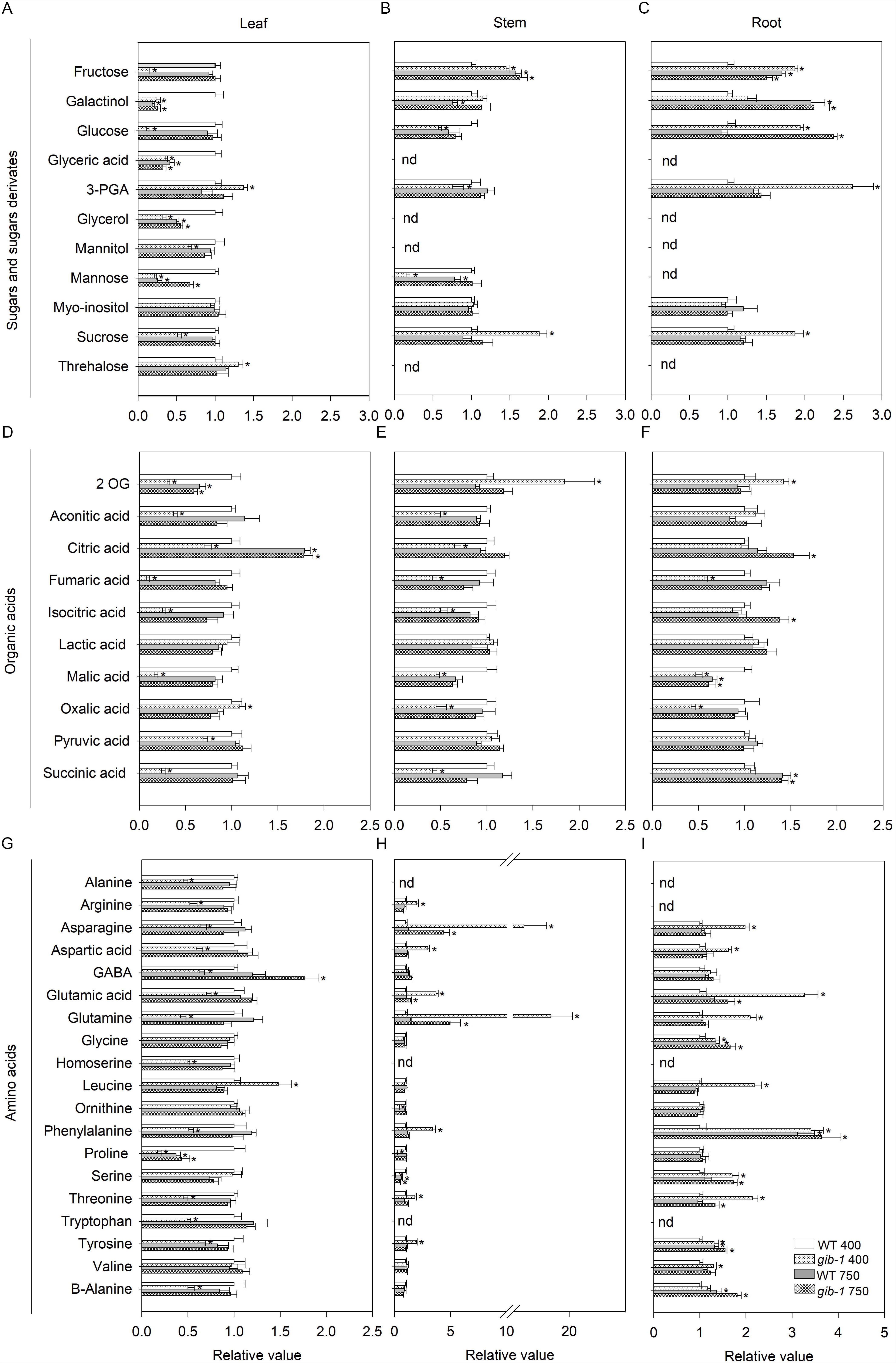
Proposed model of growth regulation by elevated [CO_2_] in tomato plants. Plants at 21 dag (a) and 35 dag (b) grown under ambient and elevated [CO_2_] and their respective meristem development stage. Elevated [CO_2_] favors photosynthesis and carbohydrate production independently of the plant age. In the juvenile phase (21 dag), gibberellin-deficient mutants can grow to the same extent as wild-type plants as a response of the cell division and expansion capacity. In the adult phase (35 dag), however, elevated [CO_2_] does not stimulate growth and gibberellin mutants show their characteristic stunted growth phenotype. This suggests that growth stimulation by [CO_2_] is highly dependent on plant developmental stage, possibly linked to the juvenile-to-adult phase transition. However, which hormones are involved in the control of these events remains an open question.

## Author contribution statement

KG conducted experiments and wrote the manuscript. LCC, performed the experiments and statistical analysis. FALB, FBC and TMP performed the experiments. WLA contributed reagents, materials, and analysis tools. DMR designed the experiments. AZ finalized manuscript writing. All authors reviewed the final version of the manuscript and approved it.

## Acknowledgements

This work was funded by a grant (443064/2014-8) from the National Council for Scientific and Technological Development (CNPq, Brazil). This study was financed in part by the Coordination for the Improvement of Higher Level Personnel (CAPES-Brazil) (Finance Code 001). We thank Joaquim Gasparini for assistance with photos.

## Supplementary material

**Fig. S1** Effects of elevated [CO_2_] on specific leaf area (SLA) and relative growth rate (RGR) of wild type (WT) and *gib-1* mutant.

**Fig. S2** Levels of nitrate, amino acids, protein and chlorophyll in wild type (WT) and *gib-1* mutant at 400 or 750 μmol CO_2_ mol^-1^.

**Fig. S3** Levels of carbohydrate in wild type (WT) and *gib-1* mutant at 400 or 750 μmol CO_2_ mol^-1^.

**Fig. S4** Levels of leaf soluble sugars in wild type (WT) and *gib-1* mutant at 400 or 750 μmol CO_2_ mol^-1^, 35 dag.

**Table S1** Relative metabolite level of leaf, stem and root from wild type (WT) and *gib-1* mutant at 400 or 750 μmol CO_2_ mol^-1^.

