## Supplementary figures and images for "Elevated CO_2_ induces age-dependent restoration of growth and metabolism in gibberellin-deficient plants"

### Supplementary Figure 1

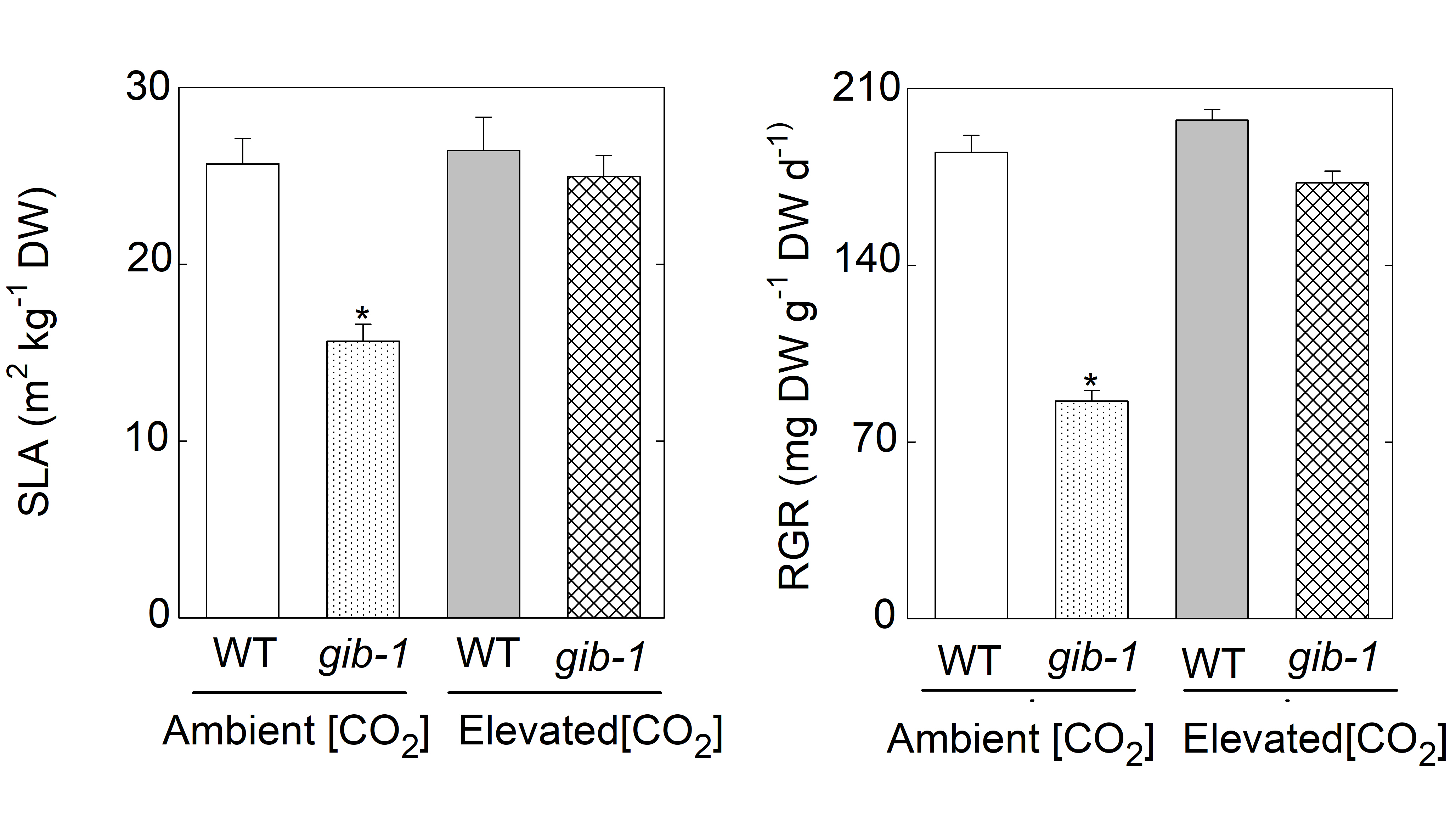

### Supplementary Figure 2

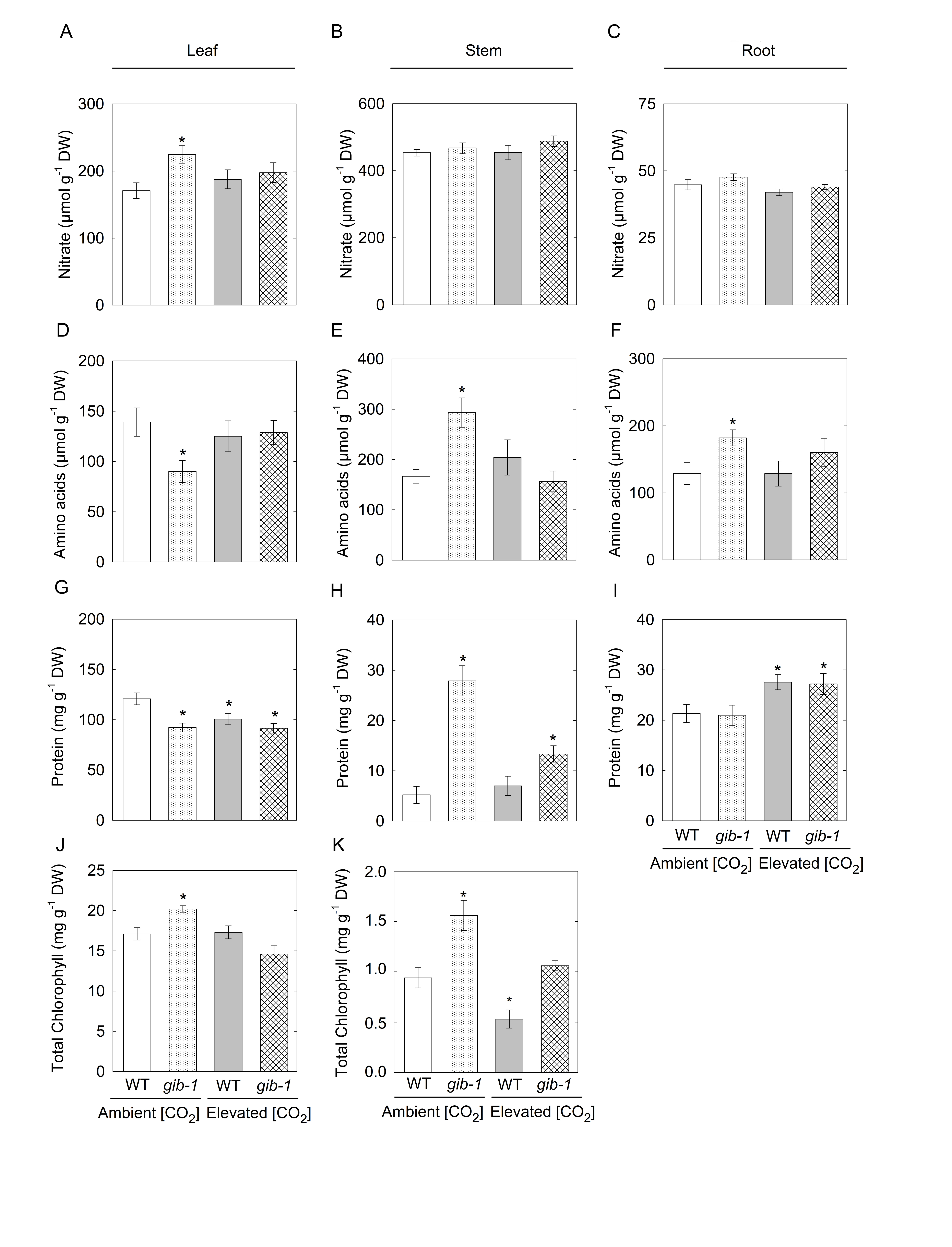

### Supplementary Figure 3

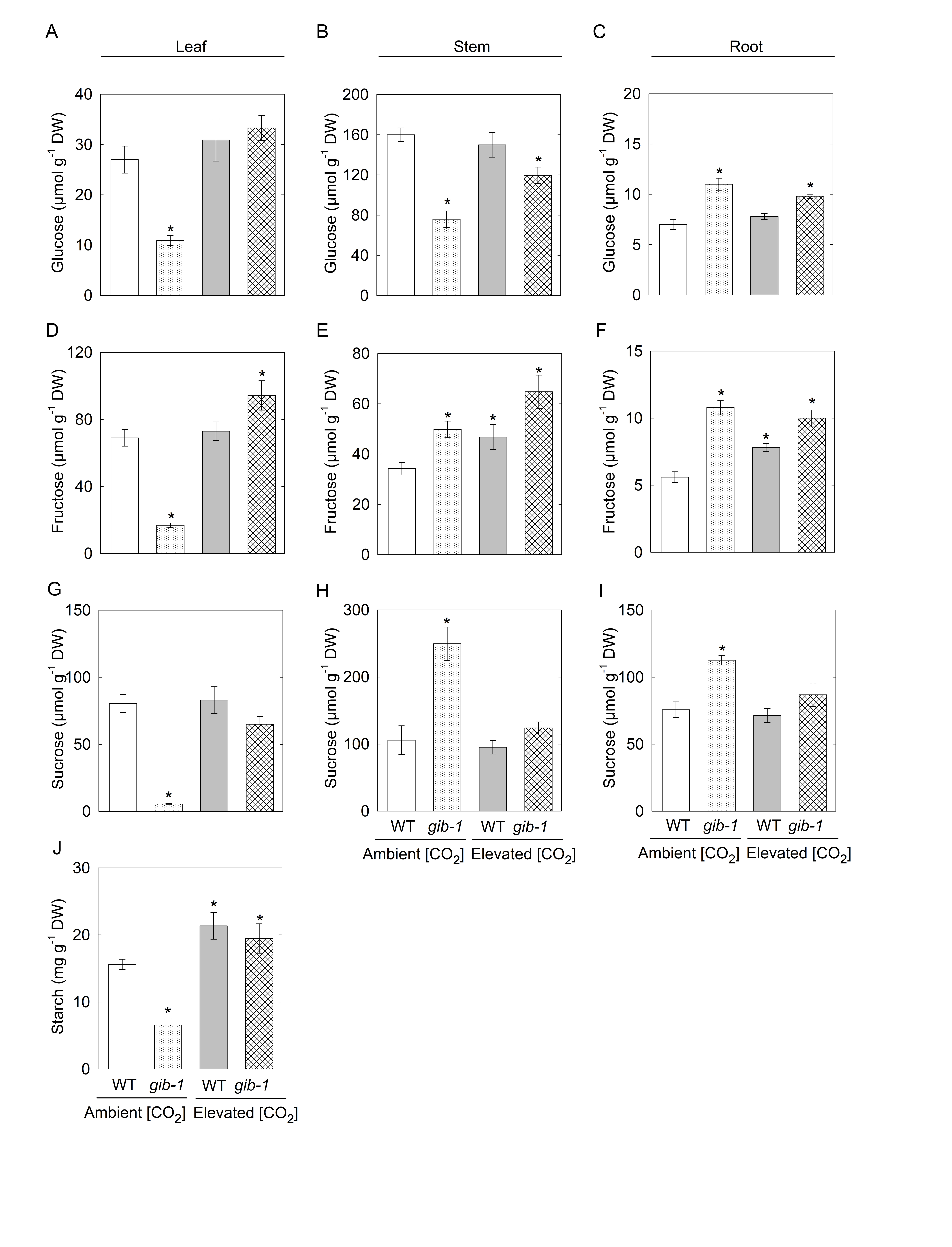

### Supplementary Figure 4

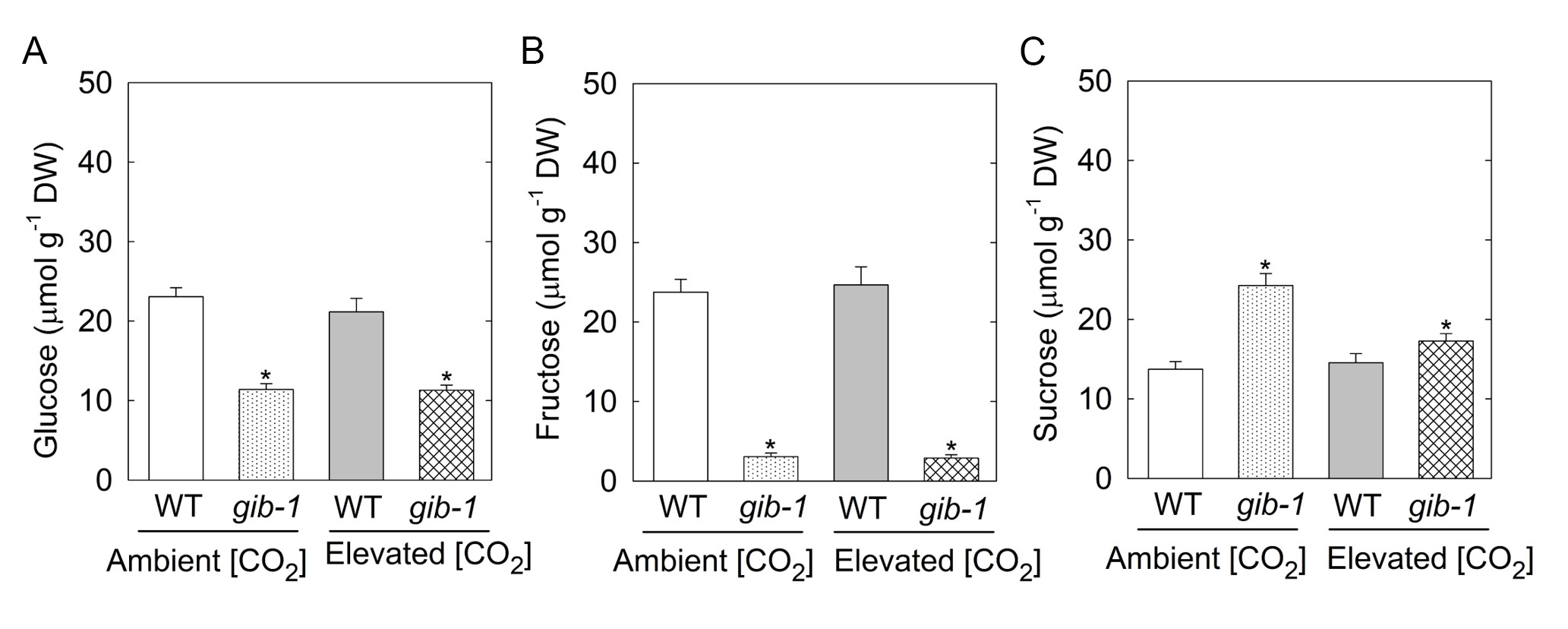
